# The buds of *Oscarella lobularis* (Porifera): a new convenient model for sponge cell and developmental biology

**DOI:** 10.1101/2020.06.23.167296

**Authors:** Rocher Caroline, Vernale Amélie, Fierro-Constaín Laura, Séjourné Nina, Chenesseau Sandrine, Marschal Christian, Le Goff Emilie, Dutilleul Morgan, Matthews Cédric, Marschal Florent, Brouilly Nicolas, Massey-Harroche Dominique, Ereskovsky Alexander, Le Bivic André, Borchiellini Carole, Renard Emmanuelle

**Affiliations:** Aix Marseille Univ, CNRS, IRD, IMBE UMR 7263, Avignon Université, Institut Méditerranéen de Biodiversité et d’Ecologie marine et continentale, Station Marine d’Endoume, Marseille, France; Aix Marseille Univ, CNRS, Institute of Developmental Biology of Marseille (IBDM), case 907, 13288, Marseille cedex 09, France; ISEM, Université de Montpellier, CNRS, IRD, EPHE, Montpellier, France; Saint-Petersburg State University, Saint-Petersburg, Russia; Koltzov Institute of Developmental Biology of Russian Academy of Sciences, Moscow, Russia

**Keywords:** morphogenesis, staining, imaging, culture, reproduction, regeneration

## Abstract

The comparative study of the four non-bilaterian phyla (Cnidaria, Placozoa, Ctenophora, Porifera) should provide insights into the origin of bilaterian traits. Except for Cnidaria, present knowledge on the cell biology and development of these animals is so far limited. Non-bilaterian models are needed to get further into cell architecture and molecular mechanisms.

Given the developmental, histological, ecological and genomic differences between the four sponge classes, we develop a new sponge model: the buds of the *Oscarella lobularis* (class Homoscleromorpha). This experimental model supplements the two other most famous sponge models *Amphimedon queenslandica* and *Ephydatia muelleri*, both belonging to the class Demospongiae.

Budding is a natural and spontaneous asexual reproduction mean, but budding can be triggered *in vitro* ensuring availability of biological material all year long. We provide a full description of buds, from their formation to their development into juveniles. Their transparency enables fluorescent and live imaging, and their abundance allows for experimental replicates. Moreover, regeneration and cell reaggregation capabilities provide interesting experimental morphogenetic contexts. The numerous techniques now mastered on these buds make it a new suitable sponge model.

**Summary statement:** Studying sponge biology is needed to understand the evolution of metazoans. We developed a new model suitable for experimental biology that allows to study morphogenetic processes with modern tools.

## Introduction

Ctenophora and Porifera are currently the two best candidates as sister group of all other extant animals (Daley and Antcliffe, 2019; Erives and Fritzsch, 2019; Laumer et al., 2019; Simion et al., 2017; Whelan et al., 2017). As a consequence, they can provide clues to understand the features held by the last common ancestor of animals, the early evolution of animal body plans and the origin of traits shared by bilaterians and cnidarians (King and Rokas, 2017; Schenkelaars et al., 2019). Nevertheless, recent comparative studies evidenced that there are still many gaps in the basic understanding of the cell and developmental biology of these animals (Dunn et al., 2015). To fill in these knowledge gaps, the development of new biological models is required.

The most recent findings in sponge transcriptomics and genomics tend to conclude that gene inventories are unable to explain the histological and morpho-anatomical differences found either between sponge classes or between sponges and other animals (Renard et al., 2018). Moreover, present knowledge in sponge cell and developmental biology mainly relies on classical, static, optic and electron microscopy. Live, physiological and functional studies remain very scarce until now (Adams et al., 2010; Borisenko et al., 2015; Ereskovsky et al., 2015; Lapébie et al., 2009; Leys et al., 2011; Nickel, 2010; Rivera et al., 2011; Schenkelaars et al., 2016; Lavrov et al., 2018). Physiological and functional data are therefore needed to finely describe sponge cell characteristics and dynamics during morphogenetic processes.

Because of their histological, ecological, embryological differences and molecular divergence, four sponge models should ideally be developed: one for each class, namely Demospongiae, Hexactinellida, Calcarea, Homoscleromorpha (Renard et al., 2018). Currently, 13 genera and 16 sponge species are being studied at cellular and molecular levels for developmental biology purposes (Schenkelaars et al., 2019) and represent “model organisms” in the historical sense because they “are convenient to study a particular area of biology” (Russell et al., 2017). But, to date, only one sponge species fulfills nearly all requirements to be considered a *bona fide* “model organisms” in the modern sense (Russell et al., 2017), with a complete set of technical tools and resources. This is the fresh water species *Ephydatia muelleri* for which functional techniques are available (Hall et al., 2019; Rivera et al., 2011, 2013; Windsor Reid et al., 2018) and easier to farm than most marine species. To study marine sponges, we therefore recently developed a new biological resource, the bud of *Oscarella lobularis* (Schmidt, 1862) that fulfills most of the criteria of a biological model, to explore the cell biology of a homoscleromorph sponge. We believe this is a major step forward for sponge cell and developmental biology research.

Homoscleromorph sponges are of particular interest to study the evolution of epithelia as it has been already pointed (Belahbib et al., 2018; Boute et al., 1996; Ereskovsky et al., 2009, 2013b; Miller et al., 2018; Nichols et al., 2006, 2012) and the unexpected higher conservation of some bilaterian genes in homoscleromorphs in comparison to other sponge classes was more recently noticed (reviewed in Renard et al., 2018). Among Homoscleromorph sponges, *Oscarella lobularis*, the type species of the Oscarellidae family, has benefited from in depth studies of its histology, embryology and ecology (Ereskovsky, 2010; Ereskovsky et al., 2009). *O. lobularis* adults are commonly found on the rocky substrates of the north-western Mediterranean Sea. The discriminant features and phylogenetic relationships with respect to other homoscleromorph species has been previously studied (Gazave et al., 2010, 2012, 2013). This species is capable of both sexual and asexual reproduction by budding (Ereskovsky and Tokina, 2007; Ereskovsky et al., 2013a; Fierro-Constaín et al., 2017). To study cell and developmental biology of these animals we have previously developed techniques on the adult stage, available all year long in contrast to embryos: *in situ* hybridization Gazave et al., 2008), pharmacological assays (Lapébie et al., 2009) and wound-healing experiments (Ereskovsky et al., 2015). Nevertheless experimenting on adults raises several issues: 1) the high number of natural specimens to be collected, 2) the dependence upon meteorological conditions to obtain samples, 3) the non-transparency of tissues (presence of pigments of different colors (Gazave et al., 2013; Gloeckner et al., 2013)) requiring to prepare histological paraffin sections to perform observation at the cellular level, which is neither compatible with the preservation of most antigens, nor with live imaging.

The present study shows how the natural properties of the bud of *Oscarella lobularis* and the technical tools that we developed on them make them a new, convenient and suitable biological resource for getting insights into sponge cellular features and processes occurring during morphogenetic processes. These resources and technical tools make the homoscleromorph sponge *O. lobularis* a very promising model organism.

## Results

### Budding is a natural process

A two-years *in situ* monitoring realized on six adults of a population of *Oscarella lobularis* from Marseille (43.212518° N, 5.330122° E) enabled us to monitor natural reproduction events and to complete the life cycle of this species (Fig. 1) (Fierro-Constain, 2016). We observed two types of asexual reproduction: by fragmentation (Fig. 1A) and by budding (Fig. 1B).

**Fig. 1:**
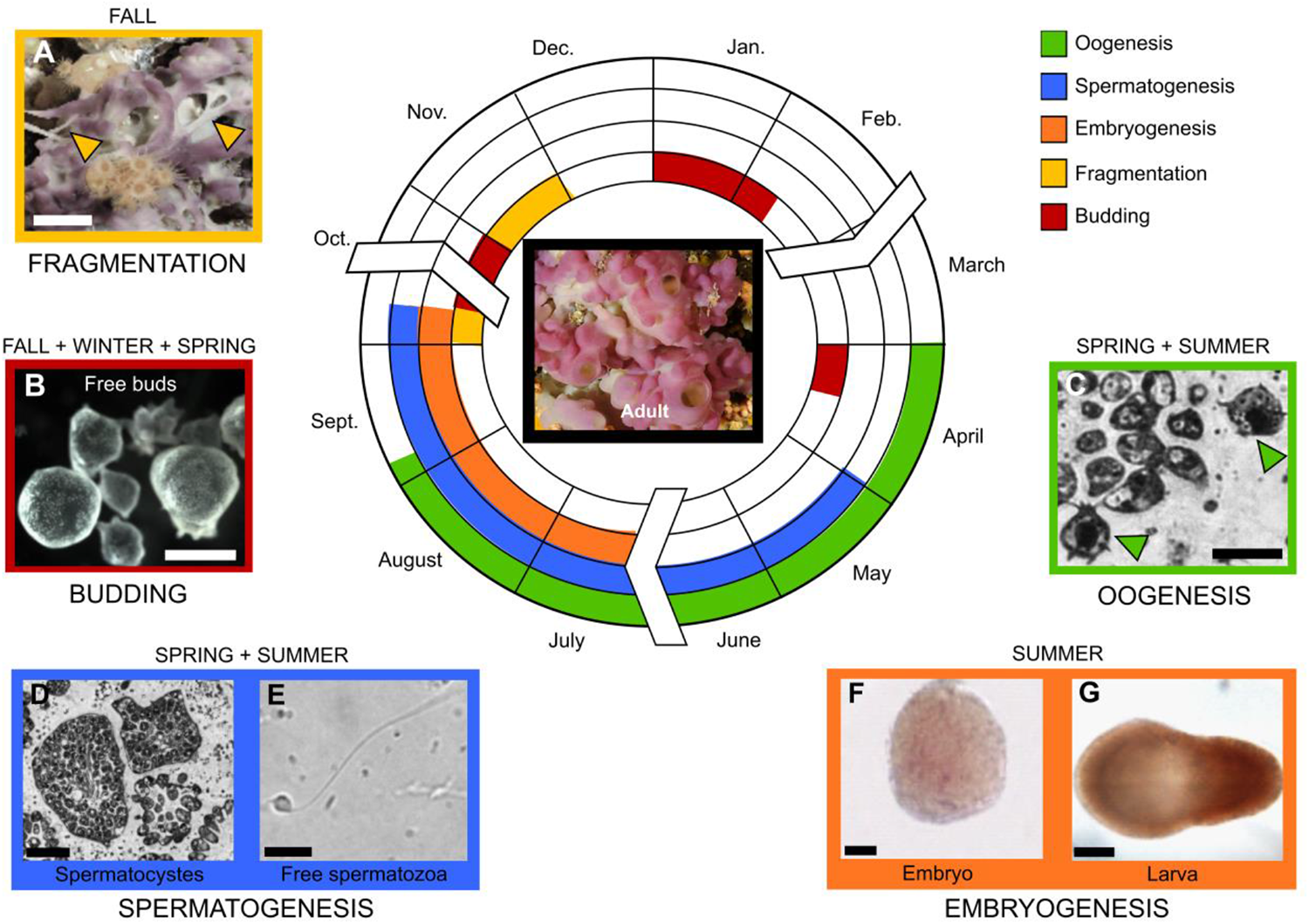
Reproduction events during the life cycle of *Oscarella lobularis*. (A) Asexual reproduction by fragmentation (arrow heads) occurs at fall, scale bar 1cm. (B) Asexual reproduction by budding yields to the release of free-floating buds, scale bar 500μm. (C-G) Sexual reproduction takes place from April to October. (C) Oocytes (green arrows) near a choanocyte chamber, scale bar 8μm. (D) Spermatozoa in spermatocysts, scale bar 25μm. (E) Released spermatozoa, scale bar 2μm. (F) Embryo dissected from maternal tissues, scale bar 50μm. (G) Swimming larva, scale bar 150μm (modified from Fierro-Constaín (2016)).

Asexual reproduction events occurred naturally several times during a year from October to April suggesting an exclusion of asexual and sexual reproduction in a same individual. Unlike sexual reproduction, fragmentation and budding do not seem to be synchronized between individuals. Budding events were observed at several periods at fall and early spring whereas fragmentation occurred at fall after the end of sexual reproduction (Fig. 1).

### Description of the budding process and its triggering in vitro

Budding adults were first observed and sampled in the sea and the terminal steps of the budding process were monitored under laboratory conditions (Ereskovsky and Tokina, 2007). Similar budding can be triggered in the lab by a mechanical stress on adults all year long: by cutting collected adults in pieces (referred hereafter as fragments).

To monitor the whole budding process *in vitro*, 3 adults of *Oscarella lobularis* were sampled. Each individual was cut into 12 fragments. The mortality caused by this mechanical stress is significant (>50%). Indeed, all the fragments that originated from one of the three individuals died, whereas 17 (out of 24) fragments from the two other individuals survived. The following description is therefore based on the monitoring of these 17 fragments.

The budding process triggered *in vitro* can be divided into three steps (Fig. 2A; Table S1) that are identical to those previously observed during natural budding (Ereskovsky and Tokina, 2007):

- *Step 1*: is characterized by a transition from a smooth body surface to an irregular surface (upper pictures of Fig. 2A).
- *Step 2:* consists in the growth of branched finger-like protrusions. All the components of the adult tissue (exopinacoderm, endopinacoderm, mesohyl and choanoderm) are concerned by this process (Ereskovsky and Tokina, 2007).
- *Step 3:* consists in the swelling of the protruding tissues via the increase of the volume of their central cavity and in the tightening of their basal pole resulting in the formation of a short stalk (Ereskovsky and Tokina 2007).

**Fig. 2:**
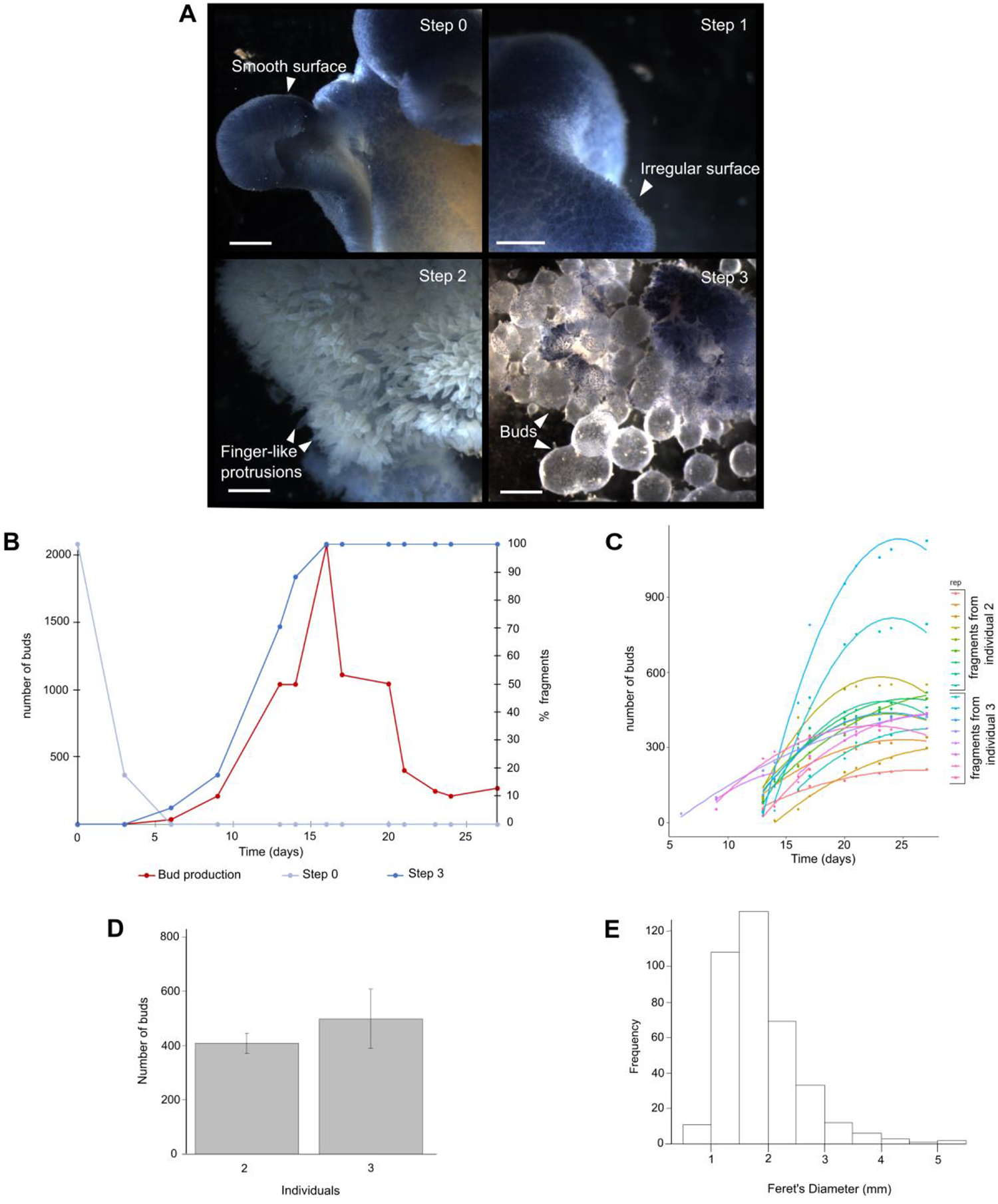
Kinetics of budding and of bud release. (A) Alive *Oscarella lobularis* fragments and buds observed using a stereo microscope; Step 0: smooth surface of a freshly sampled individual; Step 1: the surface becomes irregular; Step2: the surface is covered by finger-like protrusions; Step 3: The swelling up of protrusions results in the formation of buds that are subsequently released; Scale bars: 1mm. (B) Kinetics of budding (Table S1), budding starts 6 days after cutting and reaches a peak of bud release between 13th and 16th days after the beginning of budding process (red curve). The bud production is correlated with the percentage of fragments that reached the third step of the budding process (blue curve). (C) Daily release (cumulative) of new buds per fragment (17 fragments from two individuals) (see Table S2). (D) Average bud production for each of the 2 monitored individuals (mean: 451.4, sd: 225.56). (E) Size of stage 1 buds (Feret’s Diameter (mm)); the diameter of 84% of the bud’s ranges from 1 and 2.5 mm (mean: 1.87 mm, sd: 0.696 mm).

Time-lapse imaging of the budding process (Movie 1) shows that 4 days after cutting, fragments continue to contract rhythmically as adults do (Nickel, 2010). Budding begins on the most external part of the fragment and move to the center of the fragment (centripetal progression) until only a shred of tissue remains.

The release of free buds starts 6 days after cutting and reaches a peak between 13th and 16th days after the beginning of budding process (Fig. 2B, Table S1). The efficiency of budding differs from one individual to another (Fig. 2C, Table S2), but one cm^3^ fragment of adult can produce a mean of 450 buds (Fig. 2D).

### Bud morpho-anatomical features and development: from bud release to settled juvenile

Once free, the development of the bud can be subdivided into 4 morphologically distinct developmental stages (Fig. 3). The timing of occurrence of each stage is indicated in days since bud release (timing observed in standardized conditions). In other conditions (volume of natural sea water (NSW), frequency of sea water renewing, number of buds per well…) this developmental timing might be slightly different.

- *Stage 1 – just released spherical buds – Day 0*. At this stage, buds have a spherical shape. The central cavity is larger than it was before the release and larger than at next stages. This size ranges from 1 mm to 2.5 mm with a mean size of 1.87 mm (Fig. 2E). This difference of sizes can be explained both by a difference in the volume of the central cavity (diameter ranging from 160 to 700 μm) and by different levels of contraction between individuals (Movie 2). Indeed, buds keep on contracting rhythmically after their release from the adult tissue. These contractions are of different amplitude (Table S3): During a 105-min period, some buds can undergo a release up to 113% magnitude (mean: 53%, sd: 0,39). The general tissue organization of the bud found around the central cavity is the same as in adult despite a simpler organization of the aquiferous system (Fig. 3A). The exopinacoderm forms the most external layer of the bud and is perforated by ostia (inhalant pores). The endopinacoderm lines the canals of the aquiferous system and the internal cavity. The choanoderm is composed of choanocyte chambers communicating with exhalant canals by apopyles. The chamber dimensions range from 36 to 52 μm (mean 44,7 μm, Table S4), a size range comparable to the smaller chambers observed in adults (Boury-Esnault et al., 1984). Between these epithelial layers, the mesohyl, a mesenchymal layer, includes different cell types (next section) and symbiotic bacteria (the most abundant one being *Rhodobacter lobularis* (Jourda et al., 2015)).
- *Stage 2 – buds with outgrowths – Day 1 to Day 7*. The histological organization at stage 2 is similar to that observed at the first stage (Fig. 3B). The global shape becomes variable and asymmetric because of the formation of numerous external outgrowths. Outgrowths are only composed of exopinacoderm and mesohyl. At their distal pole, roundish cells (absent or rare at stage 1) become obvious (Fig. 3B).
- *Stage 3 – buds with an osculum - Day 8 to Day 29*. Stage 3 is characterized by the development of a polarized organization predating the basal-apical polarity of stage 4. On one pole (the future basal pole) the outgrowths are clustered, while at the opposite pole (the future apical pole), choanocyte chambers are clustered and an osculum develops (Fig. 3C). Larger choanocyte chambers are observed compared to stage 1 (up to 70μm large, Table S4, Fig. S1).
- *Stage 4 – juveniles settled on the substrate - from 1 month old*. At stage 4 (Fig. 3D), buds attach to the substrate with the osculum pointing to the surface at the apical pole. The adhesion to the substrate is probably performed by the abundant mucus present at the distal parts of the outgrowths (Fig. 4B,F). These settled juveniles (with a reduced internal cavity or atrium) are similar to juveniles formed after larval metamorphosis (Ereskovsky and Tokina, 2007; Ereskovsky et al., 2007). Some stage 3 buds fail to settle, the unsettled buds can nevertheless stay alive for several months in culture plates.

**Fig. 3:**
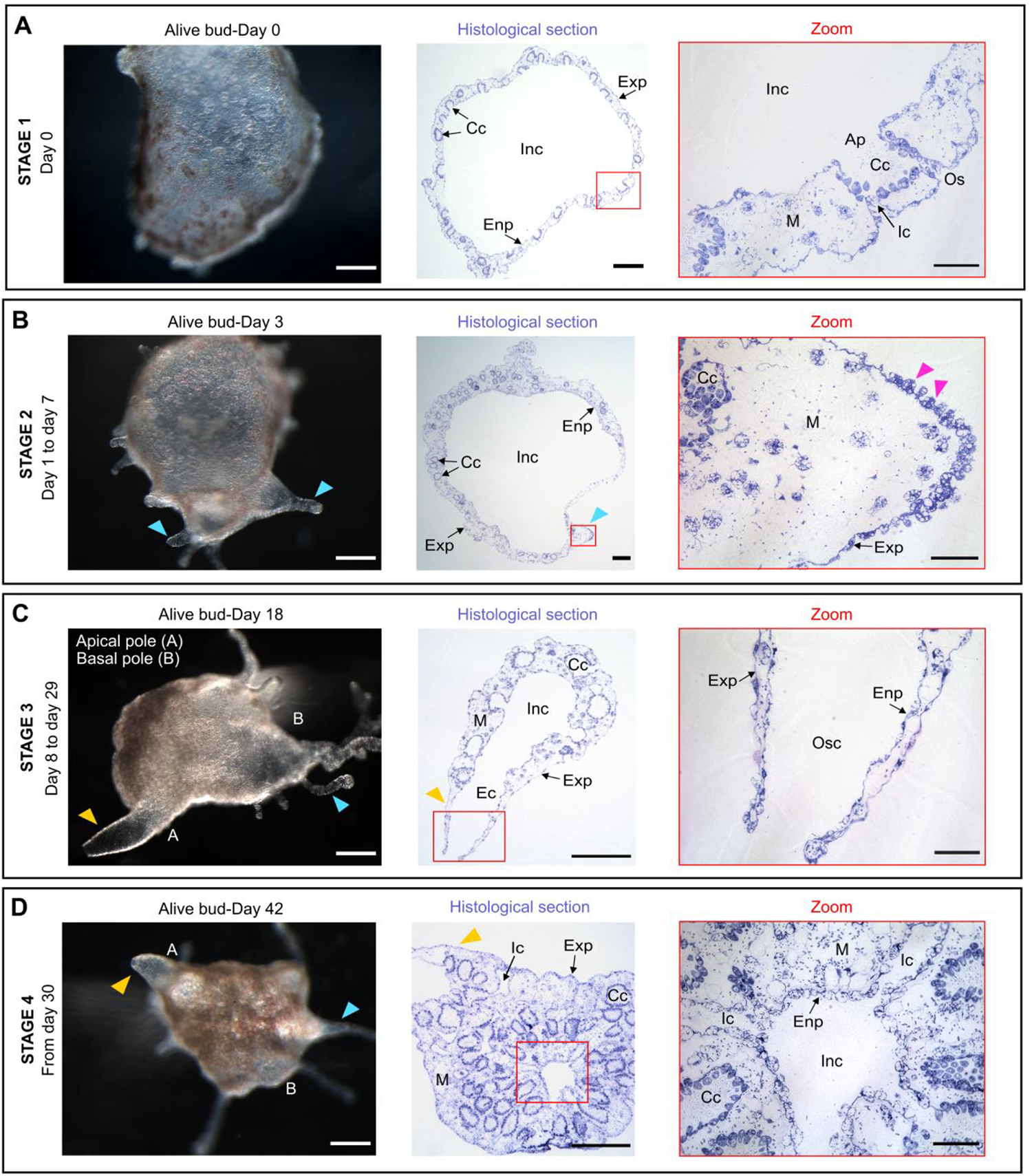
Development of bud. (A) Stage 1: Spherical just-released Bud. (B) Stage 2: Bud with outgrowths (blue arrows) devoid of choanocyte chambers (zoom in) and with distal roundish cells (pink arrows). (C) Stage 3: bud with osculum (yellow arrows) and outgrowths (blue arrows) clustered on the opposite pole. (D) Stage 4: The juvenile in settled thanks to basopinacocytes and has a reduced internal cavity (atrium). Scale bars: 200μm (first column), 100μm (middle column), 20μm (zoom). Ap: apopyle; Cc: choanocyte chamber; Ec: exhalant canal; Enp: endopinacoderm; Exp: exopinacoderm; Ic: inhalant canal; Inc: Internal cavity; M: mesohyl; Osc: osculum; Os: ostium.

**Fig. 4:**
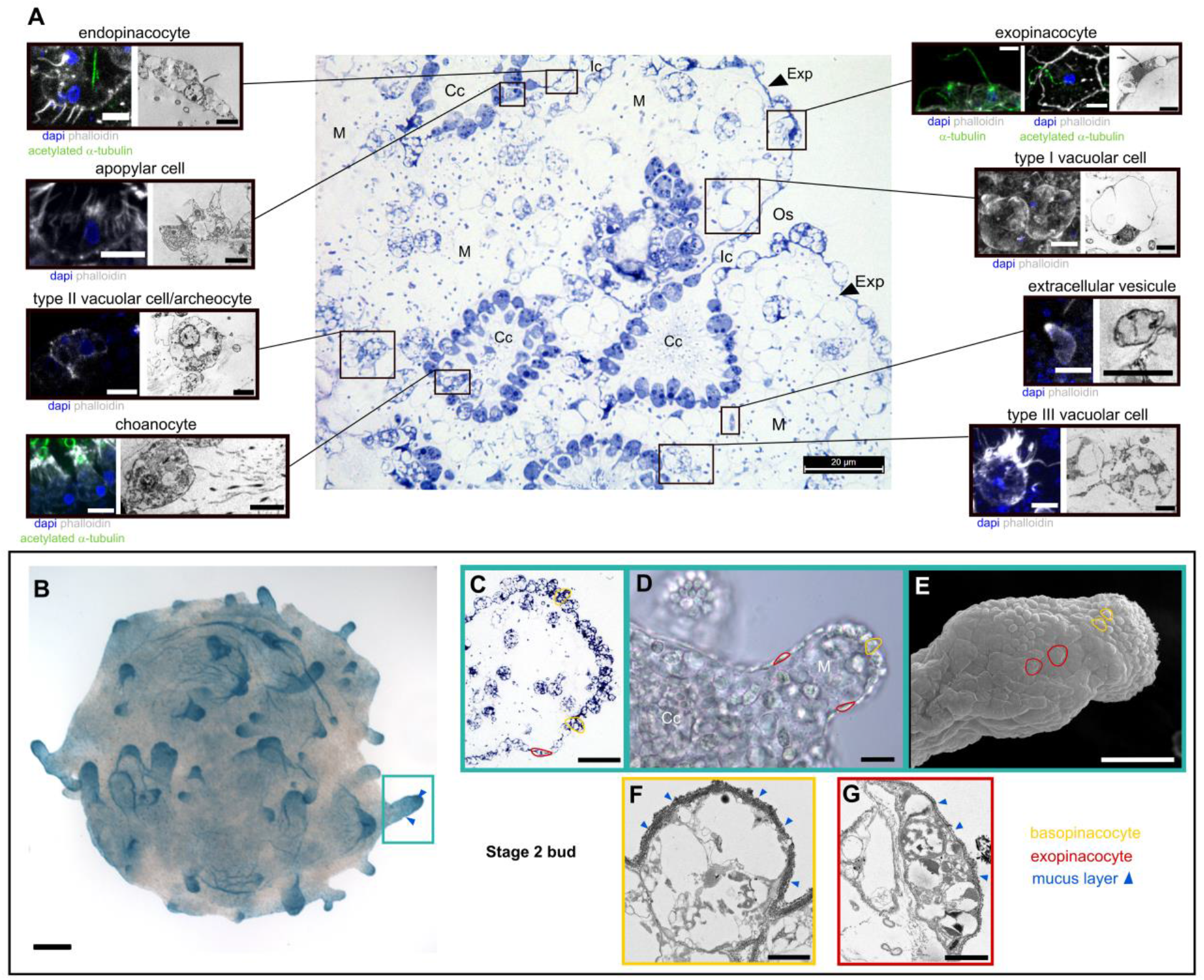
Cell types of *Oscarella lobularis* buds. (A) Position of each cell types in the tissues observed on histological sections. For each cell type, the left picture is a max intensity projection of confocal Z-stack, scale bar: 5 μm; on the right a TEM picture, scale bar: 2,5 μm. (B) External view of a stage 2 bud showing accumulation of mucus (Alcian blue staining) at the distal part of the outgrowths (blue arrows). (C, D, E) Observations of stage 2 bud outgrowth on (C) semi-thin histological section, (D) white light microscopy and (E) MEB, showing the difference between flat exopinacocytes and roundish basopinacocytes. (F) Basopinacocytes (TEM) covered by thicker mucus layer (blue arrows) than exopinacocytes (TEM) (G). Scale bars: 100μm (B); 20μm (C,D); 43μm (E); 2μm (F,G). Cc: choanocyte chamber; Inc: internal cavity; Ic: inhalant canal; Enp: endopinacoderm; Exp: exopinacoderm; M: mesohyl; Os: ostium.

In brief, the buds and the juvenile obtained *in vitro* present the same morphological and histological features than buds resulting from natural budding and the juveniles resulting from sexual reproduction.

### Description of cell types

Previous histological description of buds at stage 1 (Ereskovsky and Tokina, 2007) reported that they are composed of the same 6 cell types as adults: exopinacocytes, endopinacocytes, choanocytes, apopylar cells, type I vacuolar cells and type II vacuolar/archaeocyte-like cells (Fierro-Constain et al., 2017). By using a combination of semi-thin histological sections, confocal microscopy and electron microscopy (TEM, SEM), we reassessed cell types (Figs 4, S2A-D).

Five epithelial cell types can be easily distinguished and fit the three key criteria defining epithelial cells (Leys et al., 2009; Renard et al., in press): a polarity made obvious by the presence of an apical flagellum (Figs 4A, S2B-D) and/or the abundance of basal actin-rich filopodia (Fig. S2A-C), actin-rich adherens-like junctions (Fig. S2C), and supported by a basement membrane containing type IV collagen (Figs S2B,C). All are covered externally by glycocalyx and mucus layers, the thickness of which can differ (Fig. 4 F,G). These epithelial cell types are:

- *Choanocytes:* They form choanocyte chambers. Choanocytes are composed of a cellular body, an apical collar of microvilli and a flagellum composed of acetylated alpha-tubulin (Fig. 4A) and alpha-tubulin. The basal pole of choanocytes harbors long actin-rich cytoplasmic protrusions, filopodia, that pass through the basement membrane and contact other cells (Figs S2A,B).
- *Pinacocytes:* endo-, exo- and *basopinacocytes Endopinacocytes* line inhalant and exhalant canals whereas exopinacocytes line external parts of the bud. Both have the same ovoid to flattened shape as in adults (Ereskovsky, 2010). The exopinacocytes possess one very long (up to 20 μm, Fig. 4A) flagellum, longer than that of endopinacocytes (5-10 μm). The acetylation of the alpha-tubulin composing the flagella of exopinacocytes is more heterogeneous than that of choanocytes with a spot-like distribution along the flagellum (Figs 4A, S2C,D). As for choanocytes both SEM 3D (Movie 3) and confocal imaging on whole mounted buds evidence long actin-rich expansions at the basal pole of this cell type passing through the basement membrane and often contacting other cells (Figs S2A-C). Pinacocytes surrounding the ostia often present a fringe of actin protrusions at the edge of the pore (Fig. S2D) not reported so far. Both endo- and exopinacocytes contain more vacuoles than the ones found in adults (Ereskovsky et al., 2015). During the development of bud, round-to oval-shaped cells, with large vacuoles and very thick glycocalyx and mucus layers, are visible at the distal pole of the external outgrowths (Fig. 4B-F). These cells possess both short apical and long basal actin-rich cell extensions connecting with other cell types (Movie 4). Morphologically and functionally, these cells differ from the exopinacocytes, according to their position and putative function in bud settlement, we consider these cells as basopinacocytes, currently defined as pinacocytes affixing the sponge to the substratum by external secretion of a mucous layer (Boury-Esnault and Rützler, 1997; Ereskovsky, 2010).
- *Apopylar cell:* This cell type forms the boundary of the opening of a choanocyte chamber into an exhalant canal or central cavity. They are characterized by a flat to triangular shape, a flagellum and microvilli at the apical pole (Fig. 4A).

Three mesenchymal amoeboid and motile (Movie 5) cell types can be distinguished (Fig. 4A):

- *Type I vacuolar cell:* They are characterized by the presence of 1 to 4 large electron translucent vacuoles with few short filopodia, the nucleus is localized laterally.
- *Type II vacuolar/archaeocyte-like cells:* They have asymmetric amoeboid shape and are characterized by numerous vacuoles (empty or with fibrous inclusions, few osmiophilic inclusions), and the presence of few long actin-filopodia which often contact other cells.
- *Type III vacuolar cell type:* The size of the cellular body ranges from 7 to 11μm. They harbor a clear polarity (in contrast to archaeocytes): one roundish pole and the opposite pole with numerous thin actin-rich filopodia (>5). As for archaeocytes, numerous vacuoles of different sizes are present in the cytoplasm.

In addition, we observed in the mesohyl numerous anucleate cell-like structures which size ranges from 2 to 5 μm: we interpret them as being extracellular vesicles (EVs) (Figs 4A, S2H,I) (Raposo and Stahl, 2019). EVs have various shapes spherical, ovoid or bean-shaped with an obvious actin-rich pole involved in motility capabilities. These EVs are probably microvesicles (MVs) according both to their size and because they seem to originate from budding at the basal pole of exopinacocytes (Fig. S2G).

In brief, according to the histological criteria, the buds of *Oscarella lobularis*, would harbor at least 8 cell types. Thanks to fluorescent confocal imaging, we evidenced the presence of two new cell types and of extracellular vesicles not described previously in this species neither in adults nor in buds (Ereskovsky et al., 2009; Ereskovsky and Tokina, 2007).

### Bud are physiologically active individuals

Buds (stages 1 and 2 in particular) harbor a large central cavity allowing them to float in the water column, and flagella on the surface that, if motile, can participate to its mobility.

Time laps imaging of the exopinacocytes flagella in white light, clearly shows their beating (Movie 6). We monitored the movement of fluorescent microbeads placed on the surface of the exopinacoderm (Colibri imaging) in the presence or absence of nocodazole (Fig. S3). This preliminary experiment shows that flagella beating enables the directional movement of beads at the bud surface with a speed of about 47 μm/s. After 40 min of nocodazole treatment, a well-known microtubule inhibitor, the flux and the speed of beads drastically decreased. This observation shows that the particle flux and speed are directly correlated with the flagella beating.

In adult sponges, filtration is performed by the choanocytes which enable to pump the water unidirectionally through the body and phagocyte food particles (Renard et al., 2013). In order to establish if stage 1 buds are able to filter feed despite the absence of a complete aquiferous system, we compared the entrance of fluorescent microbeads (size 0.2μm to mimic bacteria size) in the aquiferous system between Day 0 buds (stage 1, devoid of osculum) and Day 30 buds (stage 3 possessing an osculum) after 15, 30 and 60 min of incubation (Fig. 5). The localization of fluorescent microbeads in inhalant canals and choanocyte chamber cavities shows that stage 1 buds are capable of filtering (Fig. 5A). This process is fast: after only 15 min of incubation, beads were present in inhalant canals and in the cavity of choanocyte chambers, indicating that filtration started. After 30 min of incubation, beads were observed both at the level of the microvilli of choanocytes and in the cytoplasm of choanocytes (Fig. 5A), meaning they were phagocytized. However, the comparison between buds at stage 1 and 3 after 30 min incubation indicates that whereas all choanocytes contain beads in stage 3 buds, only some of them have accumulate beads in stage 1 buds (data not shown). Stage 1 buds are capable of filtration, but their filtering efficiency is lower than that of later stages. After 30 min in stage 3 buds, some mesohylar cells contain beads (Fig. 5B).

**Fig. 5:**
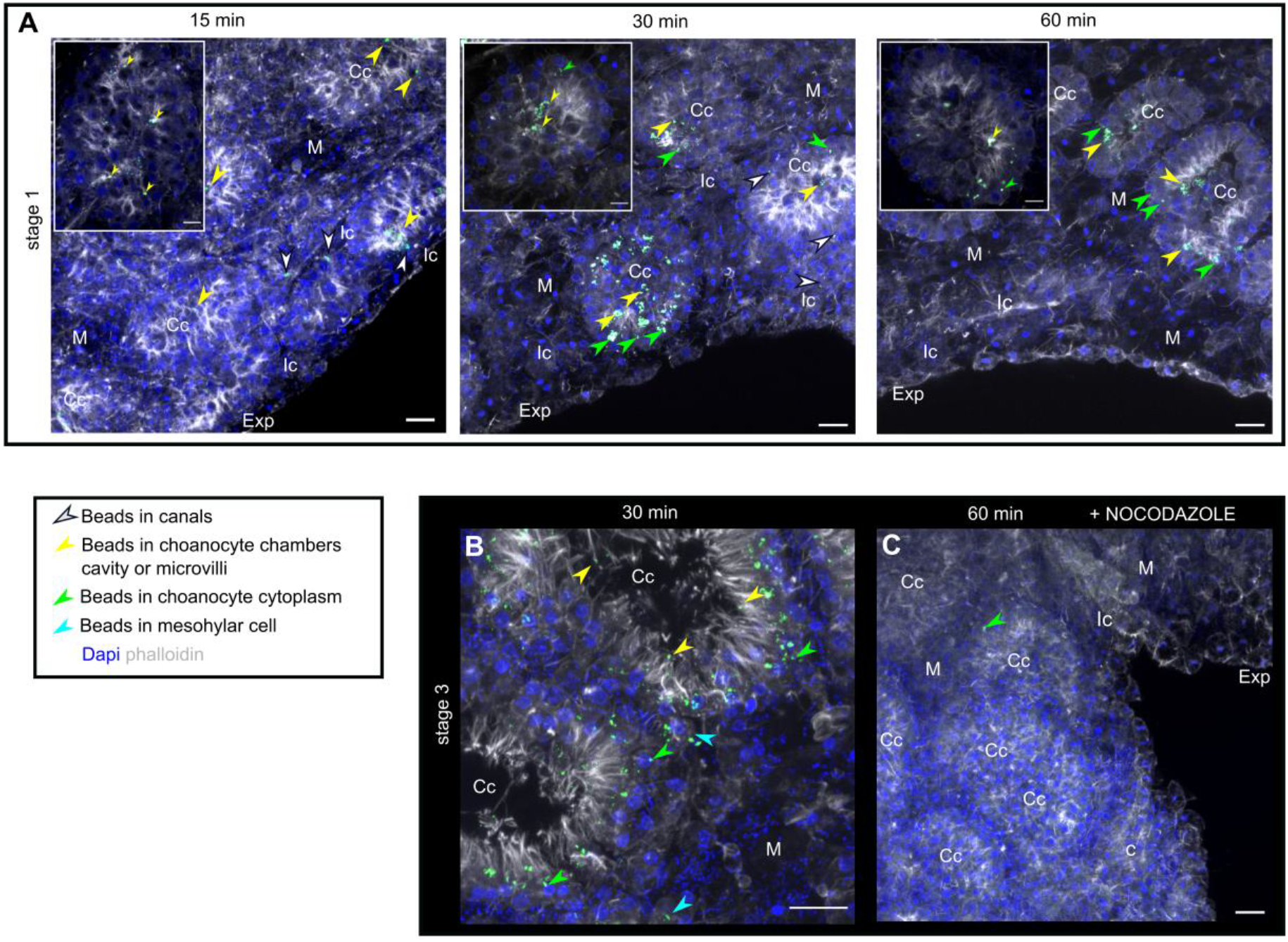
Active filtering activity of buds evidenced by 0.2 μm fluorescent beads. (A) Location of beads in stage 1 buds (devoid of osculum) after 15, 30- and 60-min incubation with beads. Arrows of different colors show different locations: canals, choanocyte chambers cavity or microvilli, choanocyte cytoplasm and mesohylar cell cytoplasm. (B) Location of beads in stage 3 buds (with osculum) after 30 min incubation with beads, in choanocytes microvilli or choanocyte cytoplasm and in mesohylar cells. (C) Absence of filtering activity after a nocodazole treatment (near absence of beads in stage 3 buds after 60 min incubation) (control: DMSO treatment, Fig. S4). Scale bar: 10 μm (A,B,C). Cc: choanocyte chamber; Exp: exopinacoderm; Ic: inhalant canal; M: mesohyl. Max intensity projection of confocal Z-stack (A, B, C).

If buds are exposed to nocodazole before incubation with beads, no or hardly no beads are found in the aquiferous system (whatever the bud stage and the incubation time with beads) (Figs 5C, S4 for control). This observation shows that the filtering activity of buds (at all stages) is an active process depending on the beating of flagella, as for adults.

### Morphogenetic processes and monitoring tools

Buds are suitable for experiments because they can be obtained in large amounts and their transparency (Movie 5) enables to stain and even track cells during morphogenetic processes. We adapted different detection and staining methods to evaluate key mechanisms (cell division, apoptosis and migration) in different developmental and non-developmental contexts.

#### Evaluation of cell renewing dynamics during bud development

To evaluate the rate of cell divisions and apoptosis events during bud development, we adapted usual approaches: Phospho-histone H3 (PHH3) immunolocalization as a marker of mitosis, 5-ethynyl-2’-deoxyuridine (EDU) cell proliferation assays and Terminal deoxynucleotidyl transferase dUTP nick end labeling (TUNEL) method for detecting DNA breaks generated during apoptosis.

According to PHH3 immunolocalization or EDU incorporation (Figs 6A,B, S5A,B) we found that about only 1% of cells divide at the same time (Fig. 6C, Tables S5,S6). Division occurs more frequently in choanocytes, and to a less extent in mesohyl cells than in other cell types (pinacocytes) (Figs 6A-C, S5A,B, Tables S5,S6). This may explain the slight increase of both the number of choanocytes per chamber and the size of choanocyte chambers between stage 1 and stage 3 (Table S4, Fig. S1).

**Fig. 6:**
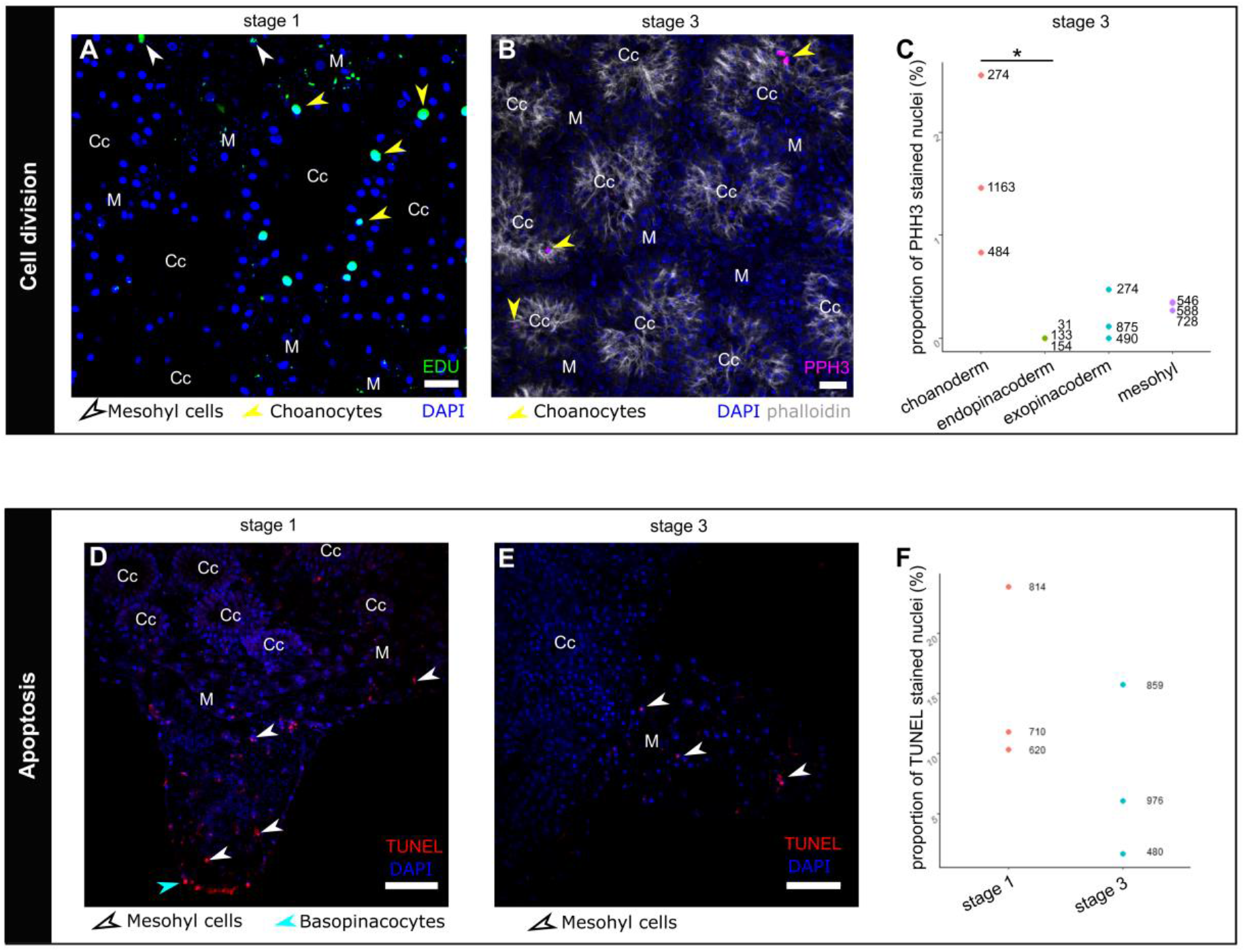
Cell renewing during bud development. (A) Cell proliferation (green: EDU) concerns mainly choanocytes (yellow arrows) and, in less extent, mesohylar cells (white arrows), at stage 1 (Table S5). (B) Cell proliferation (magenta: PHH3) also mainly concerns choanocytes at stage 3. (C) Proportion of cells observed simultaneously in division by PHH3 immunostaining at stage 3 (Table S6). Each dot represents the proportion observed for one sample, numbers indicate the total number of nuclei observed. Cell division is rarely observed in exopinacocytes (Fig. S5A,B,C) and has never been observed in endopinacocytes. The difference is significative (*) between choanocytes and endopinacocytes (Kruskall-Wallis: pvalue=0.0973; Dunn test: padj=0.0197). (D) TUNEL staining (red) at stage 1: apoptotic DNA fragmentation is mainly observed in basopinacocytes (cyan arrow) and in cells localized in the mesohyl (white arrows), (control: Fig. S5D). (E) TUNEL staining (red) at stage 3 in mesohylar cells (white arrows), in bacteria and extracellular vesicles (EVs) (Fig. S5E). (F) The proportion of cells undergoing apoptosis is not significantly different between stages 1 and 3 (values in Table S7), each dot represents the proportion observed for one sample, numbers indicate the total number of nuclei observed. Scale bars: 10μm (A, B); 30μm (D, E). Cc: choanocyte chamber; M: mesohyl. Max intensity projection of confocal Z-stack (D, E).

TUNEL assays at different stages (Fig. 6D,E, Fig. S5D for control) shows that apoptotic cells represent between 10% (stage 3) and 16% (stage 1) of all cells (Fig. 6F, Table S7). Apoptosis is heterogeneously distributed in buds and is observed more often in outgrowths. The cell types undergoing apoptosis are vacuolar cells and basopinacocytes (Fig. 6D,E). Interestingly, numerous extracellular vesicles are also stained (Fig. S5E).

These results suggest that cell renewing (division/apoptosis balance) is low and slow during bud development.

To get insights into epithelial morphogenesis, we additionally developed regenerative and dissociation-reaggregation experiments that provide two complementary non-developmental contexts.

#### Wound healing and regeneration

Wound healing capabilities of *O. lobularis* adults have been previously studied (Ereskovsky et al., 2015). Here we show that stage 3 buds (harboring an osculum) are able of both wound healing and regeneration. Stage 3 buds (n=58) were cut into two parts: one part possessing outgrowths (future basal pole of the juvenile) and one part possessing the osculum (future apical pole of the juvenile). Of the 116 half-buds 100% survived to the injury. 24 h after the injury, both basal and apical halves successfully rescued the integrity of their external epithelium (exopinacoderm), or in other words, completed wound healing. In addition, 96% of the previously recovered halves can regenerate the missing part: an apical pole with a whole differentiated osculum or a basal pole with outgrowths, in 96 h or less (Fig. 7).

**Fig. 7:**
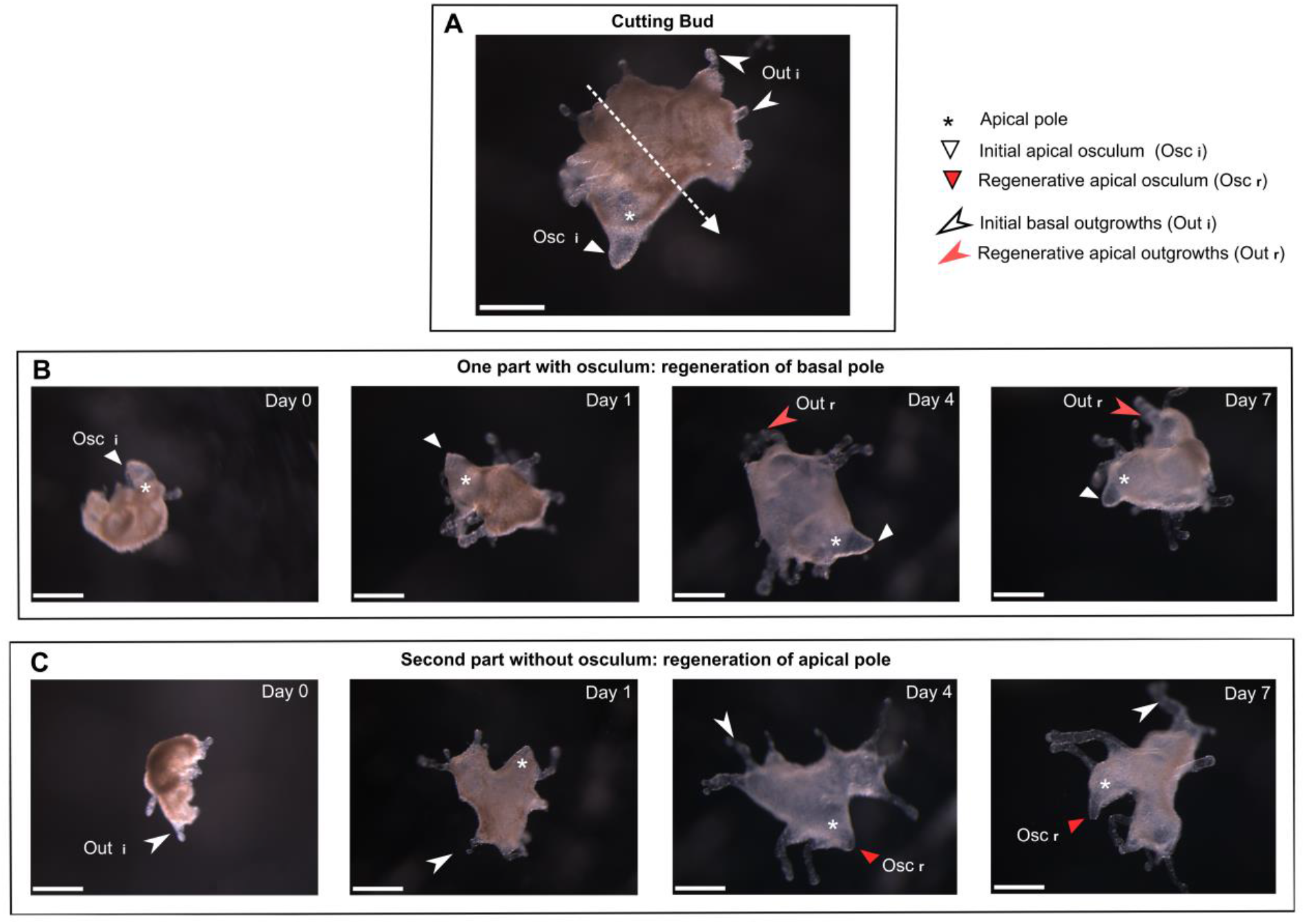
Regenerative experiments. (A) Stage 3 bud with osculum at apical pole and outgrowths at basal pole. The dotted arrow represents the section level. (B) One half with primary osculum at apical pole and devoid of basal outgrowths and (C) one half without osculum but with basal outgrowths; After 96 h, both halves regenerate the missing part: new secondary osculum or new basal outgrowth. Scale bar: 500μm (A-C).

#### Reaggregation capabilities of dissociated cell

Bud cells can be efficiently dissociated by pooling thousands of buds (from a same adult) and stirring them in Calcium-Magnesium Free Sea artificial Water (CMFSW, supplemented in EDTA) as described for other sponge (Lavrov and Kosevich, 2014). A dense cell suspension (about 0.5.10^6^ cells/mL) is thereby obtained (Fig. 8A). The viability of dissociated cells is high (>90%, data not shown).

**Fig. 8:**
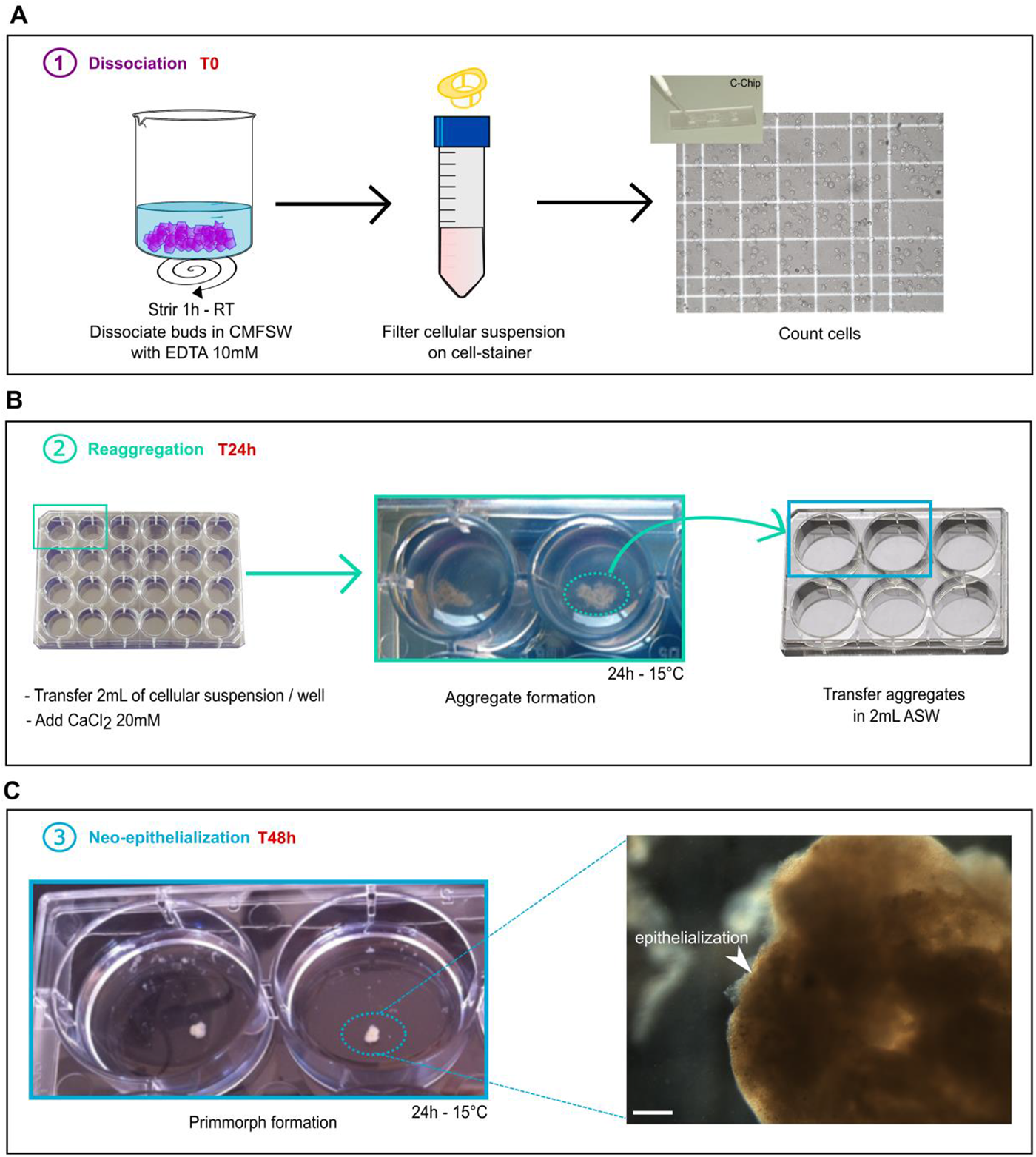
Reaggregation of dissociated cells. (A) Buds are dissociated in CMFSW and EDTA to obtain a cell suspension. (B) 24h after the addition of calcium chloride, compact aggregates are observed. (C) After 48h in ASW, Primmorph are obtained, scale bar 50μm.

The addition of calcium chlorid triggers cell reaggregation: 24 h after calcium adding, compact aggregates are obtained (Fig. 8B). Cell aggregation is followed by a neo-epithelialization: the surface of the aggregates becomes smooth. This stage is named primmorphs (Custodio et al., 1998): here, primmorphs are obtained within 48 h after calcium addition (Fig. 8C)

#### Monitoring cell migration during morphogenetic processes: Cell staining and tracking methods

To perform cell tracking during developmental and somatic morphological processes, we have developed three methods enabling to specifically stain and track choanocytes (Fig. 9):

- As previously shown, incubating buds with fluorescent beads for 30 min result in the engulfment of the beads by choanocytes. This method can be used to label choanocytes.
- As described in *Amphimedon queenslandica* (Nakanishi et al., 2014), the lipophilic marker CMDiI only stains choanocytes (Fig. 9A).
- Among different tested fluorescent labeled lectins on alive buds, two lectins, GSL1 (*Griffonia simplifolia* lectin 1) and PhaE (*Phaseolus vulgaris* Erythroagglutinin) stain specifically choanocytes, while accumulating in different sub-cellular structures (Fig. 9B,C). The long-lasting staining of choanocytes by these fluorescent lectins (up to 1 week) provides a convenient way to track these cells. For instance, using this method, we evidenced the migration of choanocytes to the mesohyl both during the natural development of bud and during preliminary regenerative experiments (Fig. 9D) Pinacocytes can also be stained by short (15-20 min) incubations with the wheat germ agglutinin (WGA, dilution 1:6000). Staining of only exopinacocytes or both endo- and exopinacocytes depends on the incubation time (Fig. 9E). The obtained staining is much less specific than the ones mentioned for choanocytes because staining of other cell types is observed quickly when increasing incubation time (>20 min) or concentration (dilution <1:6000). The use of this staining therefore requires precision and careful stain checking before undergoing tracking experiments.

**Fig. 9:**
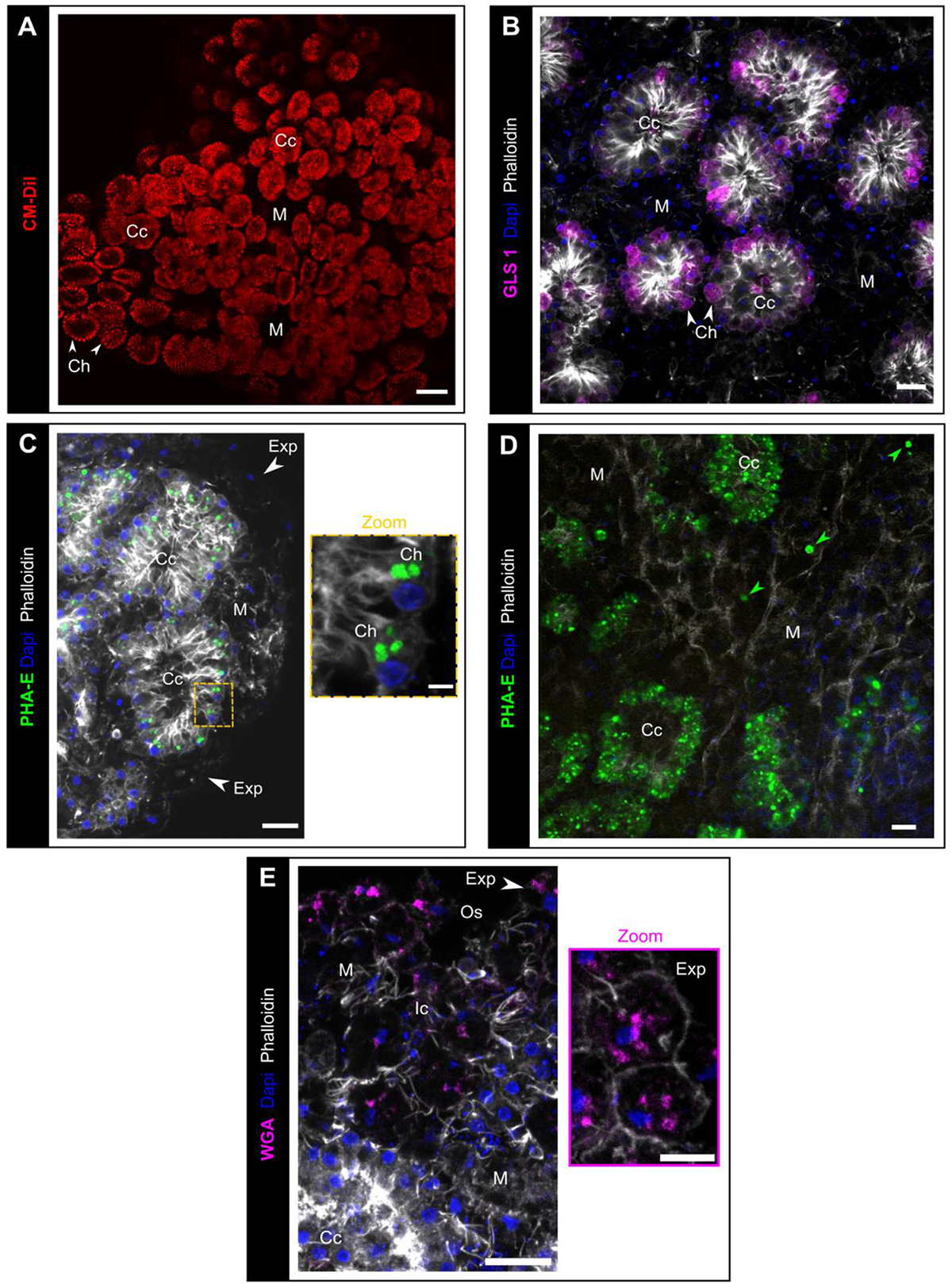
Cell staining and tracking. (A) Observation (biphotonic microscope) of choanocyte chambers in alive bud labelled by the lipidic marker CM-DiI (red). (B) GSL1 lectin (magenta) staining of choanocytes (white arrows). (C) PhaE lectin staining of choanocytes (green). (D) The presence of PhaE-stained cells in the mesohyl (green arrows) indicates a transdifferentiation from choanocytes to mesohylar cells, after 72h regeneration. (E) Rhodamine-WGA (magenta) staining of exopinacocytes (white arrow). Scale bars: 50μm (A); 10μm (B-E); 2μm (Zoom C); 5μm (Zoom E). Ap: apopyle; Cc: choanocyte chamber; Ch: choanocyte, Exp: exopinacocytes; Ic: inhalant canal; M: mesohyl; Osc: osculum. Max intensity projection of confocal Z-stack (B-E).

### Towards functional approaches

We tested several methods (electroporation, lipofectamine and mannitol-based transfection reagents and simple soacking) to transfect three types of nucleic acids (plasmid, siRNA and morpholino). For plasmid (pmaxGFP™ Vector), none of the tested methods resulted in the expression of GFP and thus we were not able to ascertain its accumulation in the cells. In contrast, fluorescent siRNA block-it and morpholino (control) molecules were successfully transfected in choanocytes by simple soacking in artificial sea water (ASW) (Fig. 10). In these two cases, neither electroporation nor the addition of mannitol improved transfection efficiency. The localization of these fluorescent synthetic nucleic acids differs: siRNA are localized in vacuoles whereas morpholinos display a more homogenous distribution in the cytoplasm. Exopinacocytes also incorporate to a less extent the morpholino (Fig. 10D).

**Fig. 10:**
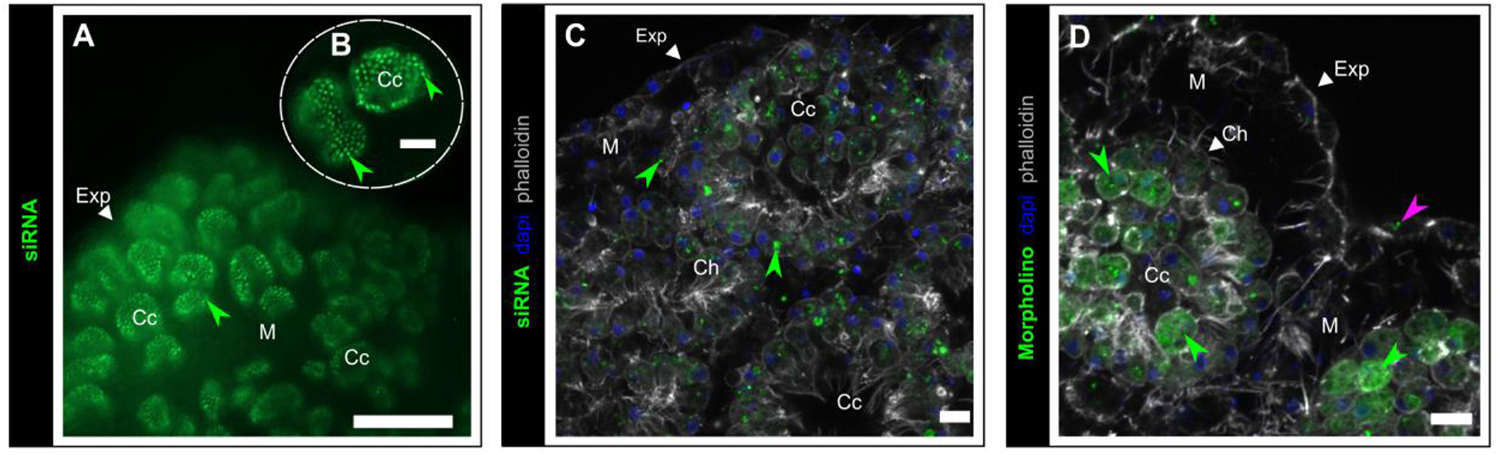
Block-it siRNA and morpholino transfection in choanocytes. (A, B) Accumulation of BLOCK-iT™ fluorescent oligo (green) (incubation 72H at 8nM) in choanocyte chambers (green arrows), (epifluorescence microscope with X20 (A) and X40 (B) objectives). (C) BLOCK-iT™ (incubation 72h at 8nM) localized in choanocyte (green arrows). (D) Standard control morpholino (green) in choanocytes and in less extent in exopinacoderm (pink arrow). Scale bars: 100μm (A), 25μm (B), 7μm (C,D). Cc: choanocyte chamber; Ch: choanocyte; Exp: exopinacoderm; M: mesohyl. Max intensity projection of confocal Z-stack (C,D).

## Discussion

### The limits of cell type definition

In the literature, according to light and electron microscopies observations, the number of cell types described in sponges ranges from 6 to at least 16 (Ereskovsky, 2010; Simpson, 1984). Our observations using fluorescent confocal imaging on whole mounted buds of *Oscarella lobularis*, enabling Z-stacking and 3D projections, yielded to the description of additional cell types in comparison to previous data on the same stage (Ereskovsky and Tokina, 2007). The distinction between cells is thus highly dependent on the technique used. A complete reevaluation of cell types with complementary tools should therefore be launched in different sponge species.

Moreover, the distinction between cell types appears in part subjective. For instance, despite a developed actin fringe on the cells surrounding the ostia (Fig. S2D), in contrast to other pinacocytes, these cells have never been considered as a distinct cell type.

Finally, considering the disparity of terminology sometimes used for similar cells, the homology of these cell types between different sponge species is unclear.

It is therefore obvious that sponge cell biology expects much of the development of single cell transcriptomic approaches, enabling to define cell types on the basis of shared regulatory networks (Arendt et al., 2016). Such data was recently acquired for two demosponge species: *Amphimedon queenslandica* (Sebé-Pedrós et al., 2018; Sogabe et al., 2019; Wong et al., 2019) and *Spongilla lacustris* (Musser et al., 2019). These first results tell us more about cell functions in sponges and evidence different sub populations in a cell type (example choanocytes) enabling to discriminate more cell types (clusters) than previously thought. We guess that the present increased number of cell types recognized in *O. lobularis* buds (8-9 instead of 6) will soon be made obsolete by ongoing single cell transcriptomic approaches.

### Two potential means of cell-cell communication in sponges

The capability of sponges to react to external chemical or mechanical stimuli has been thoroughly documented (Elliott and Leys, 2007; Leys, 2015; Leys and Meech, 2006; Leys et al., 2019; Mah and Leys, 2017; Maldonado, 2006; Nickel, 2010; Renard et al., 2009). A few sensory cells were identified: such as light sensing pigmented cells of larvae (Collin et al., 2010; Leys and Degnan, 2001; Leys et al., 2002; Maldonado et al., 2003) or mechanical-sensory ciliated cells on adult osculum (Ludeman et al., 2014). A few effector cells were also identified such as pinacocytes and apopylar cells (Elliott and Leys, 2010; Hammel and Nickel, 2014; Ludeman et al., 2014; Nickel et al., 2011). In contrast, the mechanisms of cell-cell communication and signal transduction remain unclear though electric and chemical transduction were described: action potentials in Hexactinellida (Leys et al., 1999) and neurotransmitters in demosponges (Elliott and Leys, 2010; Ellwanger and Nickel, 2006; Ellwanger et al., 2007).

Here, we observed for the first time in the homoscleromorph class the impressive length that filopodia can reach, resulting in the establishment of a real network between pinacoderm, mesohylar cells and choanoderm. These filopodia are likely to be a key element of cell-cell communication in these sponges. The significance of such cell-cell contacts was also very recently observed in *Spongilla lacustris* (Musser et al., 2019). It is possible that the existence of such a developed cell-cell interacting network was underestimated because of the common use of sections to study sponge anatomy.

Moreover, we observed microvesicles (MVs), a subtype of extracellular vesicles (EVs) (Raposo and Stoorvogel, 2013) in the mesohyl of *O. lobularis* buds. Though similar structures were previously observed in other sponge species, they were not identified as EVs: “bubbles” formed by cell shedding in another homoscleromorph sponge (Ereskovsky et al., 2007) supposed to regulate cell shape and size; and “small anucleate nurse cells” involved in the oogenesis of *Chondrilla* (Maldonado et al., 2005). The understanding of the significance of EVs in cell-cell communication in Eukaryotes including animals is recent (Mulcahy et al., 2014). It is now obvious that such microvesicles can be used to carry nucleic acids (DNA, miRNA), proteins (transcription factors) from one cell to another, can participate to immune response (antigen presentation, cell-symbiont interaction), and to apoptosis mechanisms… (Latifkar et al., 2019; Margolis and Sadovsky, 2019; Niel et al., 2018; Raposo and Stahl, 2019). Here, the TUNEL staining of some of the MVs suggests that at least some of them are apoptotic extracellular vesicles (ApoEVs), also called apoptotic bodies, supposed to play a central role in immune modulation by dying cells (Caruso and Poon, 2018). Additional studies are now needed to confirm the nature of these vesicles and evaluate their roles in cell-cell communication in sponges.

Our findings therefore represent an important step forward for the understanding of cell-cell communication mechanisms in sponges.

### Significant prospects for morphogenetic studies in sponges

The capability of dissociated sponge cells to reaggregate has been reported decades ago (Lavrov and Kosevich, 2014, 2016; Simpson, 1984).

As far as we know, dissociation-reaggregation has not been evidenced so far in homoscleromorph species (Grice et al., 2017). This study is thereby the first report of successful dissociation-reaggregation experiments in this class. This process offers a convenient experimental context to understand the molecular and cellular mechanisms involved in self/non-self-recognition (Grice et al., 2017) and in cell adhesion. Because Homoscleromorpha is the only sponge class with obvious basement membrane and adherens-like junctions, this simple experimental set-up provides the opportunity to study the dynamics of formation of these key structures of animal epithelia in early metazoans.

Moreover, the resulting primmorph can be considered as a 3D cell-culture system (Rady et al., 2019), which can be useful considering the difficulties to establish sponge cell cultures (Conkling et al., 2019).

The second somatic morphogenetic process reported here in *O. lobularis* is regeneration that offers the possibility of regenerative experiments. Most experiments performed on sponges (including the adults of *O. lobularis*, Ereskovsky et al., 2015; Fierro-Constaín et al., 2017) are wound-healing experiments instead of regenerative experiments (Lavrov et al., 2018). Indeed, the authors focus on the restoration of exopinacoderm integrity. Here we performed regenerative studies on buds by cutting off a key functional structure of sponges: the osculum. It is well-known that wound-healing and regenerative capabilities are highly variable from one animal phylum to another. Therefore, the origin and evolution of these properties (homologous or homoplasic features) is still debated (Lai and Aboobaker, 2018). The reproducible and quick osculum regeneration in buds of *O. lobularis* provides a simple and convenient model to study cellular and molecular mechanisms involved during sponge regeneration.

### Launching knock down assays

The last decade of evo-devo studies on sponges was dedicated mostly to the acquisition of transcriptomic and genomic data. These significant data have highlighted the conservation of most of the main genes involved in developmental processes of the other metazoans. The next step is to understand how similar genetic toolboxes can result in dissimilar body organization and features. To answer this question, three main axes of research have to be undertaken: increasing gene expression studies; reinterpreting sponge cell biology (previous section); and developing knock down protocols. Interfering with gene expression is required to study gene functions, but it undoubtedly the biggest challenge for sponge biology (Lanna, 2015; Schenkelaars et al., 2019).

So far, efficient and reproducible transfection and interference was only used in the freshwater demosponge *Ephydatia muelleri* (Hall et al., 2019; Rivera et al., 2011, 2013; Windsor Reid et al., 2018). Four other papers report gene transfection assays in other sponges: one in another freshwater species *Spongilla lacustris* and three in marine demosponge species (Grasela et al., 2012; Pfannkuchen and Brümmer, 2009; Revilla-I-Domingo et al., 2018; Rivera et al., 2011). Unfortunately, transfections either failed or their efficiency was too low to perform reliable knock-in or knock-down. One of the difficulties is probably that most marine sponges tolerate only low salinity changes. Altering the salinity of media (for RNA stability or electroporation purposes for example) remains a major challenge for the future.

Here, we report the first tests of transfection in a homoscleromorph species, even if many conditions (types of plasmids, promoters, transfection reagents, choice of targets.) remain to be tested. The efficient and reproducible incorporation of fluorescent-standard control morpholino in the cytoplasm of choanocytes (with a homogenous distribution) is a first encouraging step. Indeed, these synthetic nucleic acids are well known for their high stability and have already been used successfully in marine species (Heasman, 2002). Nonetheless, at this point, we cannot presuppose the ability of these transfected molecules to efficiently interfere with gene expression or translation in this species.

### Oscarella lobularis reaches the top 3 of sponge “models”

The embryology, life cycle, development, regeneration, histology of the homoscleromorph sponge *Oscarella lobularis* are well described. Moreover, its genome annotation and single cell transcriptomic are in progress to complete this knowledge (Belahbib et al., 2018; Jourda et al., 2015). Like other species, *O. lobularis* reproduces both sexually and asexually. We managed to artificially induce in the lab a natural asexual reproduction event: budding, that occurs several times a year in natural conditions.

The buds obtained from *O. lobularis* are numerous and easy to maintain several months in the laboratory in natural sea water or up to one week in artificial sea water and can travel for up to 10h. These buds therefore offer the possibility of performing a wide range of assays (on individuals either genetically identical or not). Buds are small so that incubation in small volumes is possible (100 μL) which is convenient for expensive or toxic reagents. Moreover, in contrast to gemmules, a dormant stage produced in freshwater sponges (Simpson, 1984), *Oscarella* buds are physiologically active and can be used for physiological assays.

Buds are transparent therefore enabling fluorescent imaging on both fixed and live specimens. The present paper is also the first report of the use of fluorescent lectins for specific cell labeling in sponges. This is a significant step forward for cell tracking experiments to evaluate transdifferentiation events. We now also master usual tools to study cell migration, cell proliferation or apoptosis during the natural development of buds but also during regeneration and dissociation-reaggregation experiments.

Altogether, these technical progresses in *Oscarella lobularis* buds provide new tools to better understand sponge cell biology and the cellular and molecular mechanisms involved in this sponge class. The variety of experimental tools available will establish *O. lobularis* buds as a biological model comparable to that of the juveniles of the demosponge *Amphimedon queenslandica*, and the freshwater species *Ephydatia muelleri* (table 1).

**Table 1:**
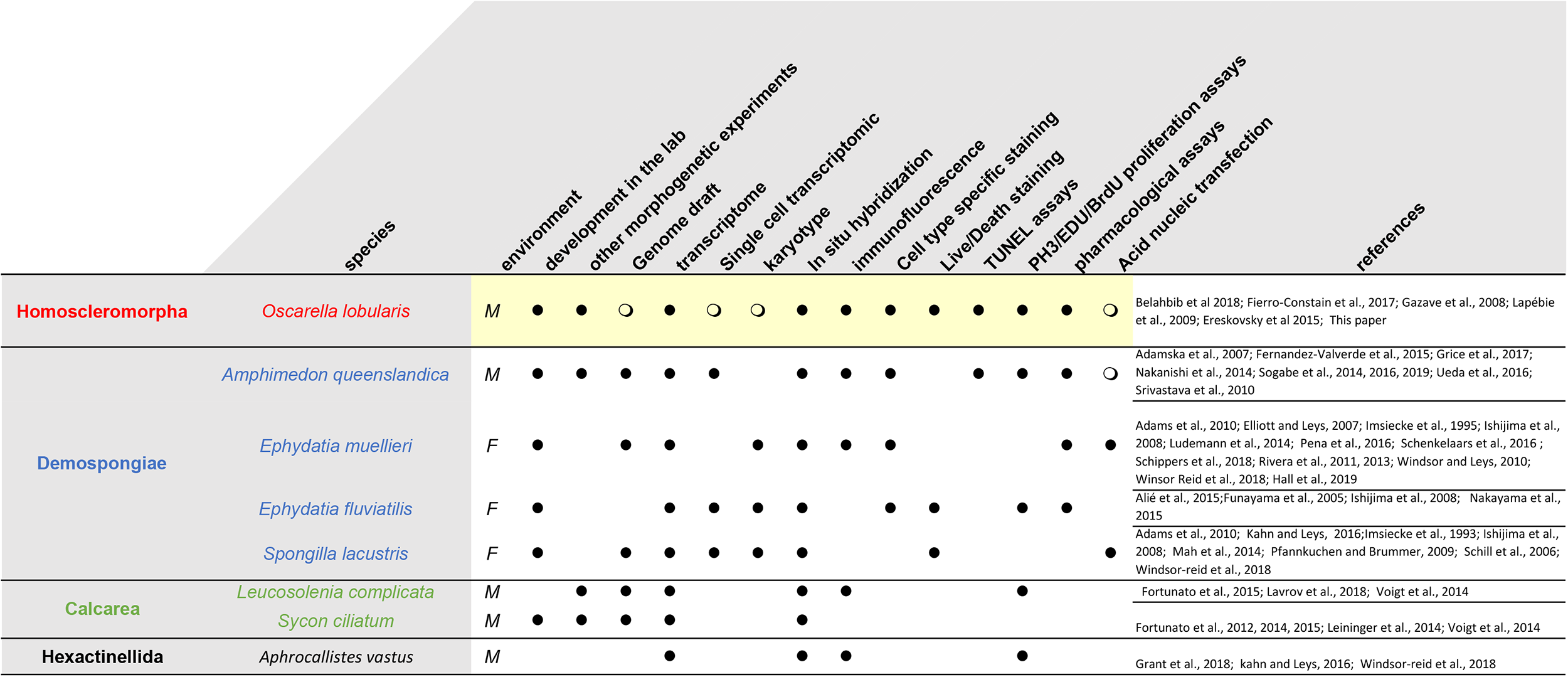
Data and tools available for the main sponge species used for evo-devo studies (for review Schenkelaars et al., 2019)

Though sponges have been studied for over a century, the main sponge “models” (Lanna, 2015; Russell et al., 2017) according to the availability of data and techniques belong to the demosponges. The emergence of a new biological resource pertaining to a distinct class, i.e. the buds of *Oscarella lobularis* (Homoscleromorpha), is highly significant for evo-devo comparative approaches (Renard et al., 2018). Moreover, because of the features of homoscleromoph epithelia compared to that of the other sponge classes, this model is of particular interest to study the evolution of epithelia and epithelial morphogenesis.

## Material and methods

### In situ life cycle monitoring

The *in situ* monitoring of a small population in the bay of Marseille (western Mediterranean Sea) was carried out. Six adults of different sizes and colors located on the north edge of Maïre Island (43.212518 ° N, 5.330122 ° E), in a large, shallow and rocky cavity (6-10 m). The photographic monitoring was implanted once a month from April 2014 until October 2015. In addition, a small fragment of each individual was sampled twice a month, and fixed (in 4% PFA overnight (O/N)). Histological observations and *in situ* hybridization were realized on sections of these small fragments (Fierro-Constaín et al., 2017) to determine the beginning and end of the sexual reproduction period. Sampling compliance provided by the “Préfet de la région PACA”.

### In vitro budding and maintain of buds in lab conditions

Freshly collected Adults (in the northwestern Mediterranean Sea (Marseille Bay)) are carefully cleaned under stereomicroscope to get rid of other organisms intermingled with sponge tissues (cnidarians, annelid tubes, algae). Then, samples are cut into pieces of about 1 cm^3^ with a sterile scalpel. Each fragment is placed in a well (3.5 cm diameter) of a 6 wells cell culture plate containing 8 mL of natural sea water (NSW). Culture plates are maintained at 17°C in a thermostated room, NSW is renewed each day until the end of budding.

Once buds are released from the adult fragment, they are transferred to Petri dishes (14 cm diameter) containing NSW and maintained at 17°C. NSW is renewed once a week.

To estimate the efficiency of budding *in vitro*, 3 adults of *Oscarella lobularis* sampled *in situ* in January 2018 were placed in thermostated room at 17°C overnight (ON), before starting experiment. Budding experiment was performed as described above. Each sample was cut into 12 fragments. Samples were observed and imaged, and buds counted every day. The bud production per day was analyzed using linear mixed effects models (LMM) that were implemented within lme4 package in the R software (Core Team, 2013). We considered time as fixed effects and we allowed linear and/or quadratic effects on the dependent variable. We also added individuals (with random slope of time within individuals) and replicates (fragments) nested in individuals as random effects.

After observation of the residual’s distribution of the models, we retained Gaussian distribution. We tested the significance of fixed and random effects respectively with *F* tests and likelihood ratio tests. When two models were equivalent, we retained the most parsimonious model.

### TL Photonic, SEM and TEM microscopy observations of budding adults and buds

For scanning (SEM), transmission (TEM) electron microscopy and semi-thin sections’ preparation a standard fixation method for TEM was used: glutaraldehyde 2.5 % in a mixture of 0.4M cacodylate buffer and seawater (1 vol., 4 vol., 5 vol., 1,120 mOsm) and post-fixation in 2% OsO_4_ in sea water as first described in Boury-Esnault et al. (1984). Semi-thin sections were stained with toluidine blue. For SEM the specimens were fractured in liquid nitrogen, critical-point-dried, sputter-coated with gold-palladium according to the protocol previously used for this species (Johnston and Hildemann, 1982).

### Observation of mucus on buds’ surface

Buds were fixed in 3% paraformaldehyde (PFA) in phosphate buffered saline (PBS) at 4°C ON, then washed 3 times with 1mL of PBS (15 mins each) at room temperature (RT). Buds were then incubated 10 min in 1mL of Alcian Blue solution (Alcian Blue 1% in Acetic Acid 3%, pH 2.5) (20:1000 in NSW) at RT, rinsed in 2 baths with NSW and immediately mounted with ProLong^®^ Antifade Mountant. After polymerization (40 min at 37°C in obscurity) the samples were observed with a stereomicroscope.

For TEM observations, we used a special fixation with ruthenium red (Ereskovsky, personal communication). Buds were fixed in buffered 2.5% glutaraldehyde + 0.1% ruthenium red for 2 h at 4°C, then rinsed 3 times with the 0.1M Na-Cacodylate buffer (30 min each) at RT, postfixation in buffered 1% OsO_4_ + 0.1% ruthenium red in the same buffer for 3 h at RT, then rinsed 3 times with the same buffer (30 min each) at RT, dehydration in ethanol series and inclusion in a resin.

### Immunofluorescent assays on whole bud

Buds are usually fixed (6 h to 24 h) in 3% paraformaldehyde (PFA) in PBS at 4°C. After fixation, buds were washed in three baths of PBS (15 min each), then, buds were incubated 45 mins in a blocking solution (Blocking Reagent (BR) (Roche) at 2% and saponine (Sigma) at 0.1% in PBS). Incubation with primary antibodies (against acetylated-α-tubulin (Sigma) (1:500), α tubulin (Sigma) (1:500), or type IV collagen (Eurogentec) (1:200)) was performed in a blocking solution at 4°C ON. Buds were washed in 3 baths of PBS (5 min each) at RT. Buds were incubated at RT for 1 h with Alexa Fluor-coupled secondary antibodies (1:500 in PBS) (anti-mouse or anti-rabbit (Jackson ImmunoResearch or LifeTechnologies)). After counterstaining with Alexa fluor-(488, 568 or 647) coupled-phalloidin (1:1000; Thermo Fisher Scientific or Santa Cruz Biotechnology) and DAPI at 2 mg/mL stock solution (1:500) 15 min in PBS, samples were rinsed and mounted in ProLong™ Diamond Antifade Mountant or ProLong™ Gold Antifade Mountant (Invitrogen).

Homemade antibodies against the type IV collagen of *O. lobularis* were targeted against the QTISDPGEEDPPVSKC peptide from the NC-1 domain (GWLLVVHSQTTDLPSCPDGSAMLYSGYSFLQSLGNGYGHGQDLGRPGSCLRVFSTMPFMHCSSSQCLVAERSDISYWLTTDQVMPVNQEDDDGNIAKYISRCAVCETKSRTIAVHNQSTGVPDCPRGWDSVWQGFSFIAQTNDGAEGGGQSLSSPGSCLRRFYPMPYIECYPRGNCKYSSPGLSFWLSPLDDTEPPFDPPVRQTISDPGEEDPPVSKCRVCSKR). They were raised in rabbits using the speedy protocol from Eurogentec (Seraing, Belgium) and purified against the same peptide before using for immunofluorescence at 1:200.

Imaging of tissues and cells were performed with an inverted confocal microscope Zeiss Axiovert 200M or confocal microscope Zeiss Axio Imager Z2 780 or 880 (objective X63-NA 1.4). The FIJI software (Schindelin et al., 2012) was used for images editing (crop, color balance, choice of colors, z-stacking, 3D projection).

### SEM3D observations of buds

Samples were prepared using the NCMIR protocol for SBF-SEM (Deerinck et al., 2010). Imaging was carried out on a FEI Teneo VS running in low vacuum (30 Pa), at 2kV and using a backscattered electrons detector. Acquisition voxel size was 5×5×40 nm.

### Time Laps imaging of budding, buds and cells

For the time-lapse imaging of the budding process, adult fragments in 6 well cell culture plate containing 8 mL natural sea water (NSW), were placed in a dark chamber at 17°C. We used a CANON EOS 750D camera with an EF-S 60mm 1/2.8 1:2.8 USM *Ultrasonic Macro Lens* to acquire one picture every 15 min during 6 days.

We used the same protocol for imaging the contractions of the free buds *in vitro*, in this case the time-lapse imaging was performed by taking one picture every 5 min for 48 h. For each bud, the maximum and the minimum sizes were measured in pixel to evaluate the amplitude of bud contraction/release. The mean and the standard-deviation (sd) were calculated on the R software (https://www.r-project.org/).

To observe cell motility, time-lapse imaging was performed on a stage 3 bud placed into 6 wells cell culture plate with natural sea water, at 22°C. We used an inverted microscope Zeiss Axio Observer Z1 in DIC TL, with an objective X20-NA 0.7, to acquire one picture every 10 s during 10 min (total 61 pictures).

To observe cilia beating on the exopinacoderm, a time-lapse imaging was performed on a bud with a microscope Zeiss Axio Imager Z1 with DIC II (objective X40-NA 1.3) with a rate of one picture every 34.3 ms during 30 sec.

### Testing Filtering and cilia activities with fluorescent beads

Day 0 (stage 1) and Day 30 (stage 3) 10 buds were incubated 15 min, 30 min and 1 h with 0.2μm FluoSpheres™ Carboxylate-Modified Microspheres yellow-green (505/515nm) (Molecular Probes) in 2 mL (1/10000) of NSW at 17°C. Buds were then rinsed 3 times with NSW and fixed (3% PFA) at 4°C ON. After DAPI/ Phalloidin counterstaining and mounting as described in the previous section, imaging of samples was performed with a confocal microscope Zeiss LSM 880 Leica. The same experiment was reiterated after a 40 min nocodazole (33μM) treatment (nocodazole is an inhibitor of tubulin polymerization, process required for cilia beating). As a control, buds were incubated 40 min in a DMSO solution (1/1000, the same concentration as the one used to solubilize nocodazole).

To characterize the beating of exopinacoderm cilia, buds were incubated with 0.5μm FluoSpheres™ Carboxylate-Modified Microspheres Red (580/ 605nm) in a Petri dish with a coverslip. The movement of fluorescent beads in live, was imaged with an inverted optical microscope Axio Observer Z1 with a LED system illumination (Zeiss). In order to estimate the flow created by cilia, the path of the beads was measured with Fiji plugin “Track Mate” with “DoG detector” and “LAP tracker” options. The same experiment was reiterated after a 40 min nocodazole (33μM) treatment.

### TUNEL assays

Buds were fixed 20 min with 3.7% PFA in filtered seawater and then permeabilized for 20 min at RT with 0.2% Triton X-100 in TS solution (150 mM NaCl, 25 mM Tris, pH 7.5). After rinses, TUNEL staining (Roche, *In situ* cell death rhodamin detection kit) was performed according to the manufacturer’s instructions. Nuclei were detected with DAPI labelling (Euromedex).

Samples were analyzed with a confocal microscope Leica TCS-SPE (Montpellier RIO Imaging platform, France). The number of nuclei and stained nuclei (table S6) was estimated using FIJI cell counter. A statistical analyze with R is performed using the Kruskall-Wallis test follow by a Dunn test.

### Cell proliferation assays

Cell division activity was estimated on both young (stage 1, 0-3 days old) and older buds (stage 3, 1 month old) by two different means: anti Phospho-histone H3 (PHH3) immunolocalization (Abcam ab5176) antibody (1:200) (see previous section for details on immunofluorescent assay protocol) and EDU staining. EDU staining was performed according to manufacturer recommendations (Click-iT™ and Click-iT™ Plus EDU Alexa Fluor™ 488 Imaging Kit (Invitrogen). Assays were performed with 10 μM, 50 μM and 200 μM in NSW: the optimal concentration was then fixed to 50 μM. Assays were performed in 24 well culture plates, each plate containing 1mL of 50μM EDU solution. Buds were incubated for 2, 6 or 24 h at 17°C. Only 6 h and 12 h incubation times yielded suitable results. Observations were performed with optical sectioning systems: an apotome microscope Axio Imager Z1 (EDU) or confocal microscope Zeiss Axio Imager Z2 780 (PHH3). The number of nuclei and stained nuclei (tables S4 and S5) was estimated using FIJI cell counter. A statistical analyze with R is performed using a one factor Anova.

### Regenerative experiments

Stage 3 buds (n=58, 3 independent technical replicates), with a differentiated osculum, were cut into two halves: one half corresponding to the future basal pole of the juvenile, and one half possessing the osculum and corresponding to the future apical pole of the juvenile (Fig.7). The 116 half-buds obtained after cutting) are placed in 24-wells culture plates containing 2mL of Natural Sea Water (NSW, renewed every day), and the culture plate placed at 17°C. Wound healing and regeneration were monitored once a day during 96 h.

Two additional preliminary experiments consisted in staining choanocytes with the PhaE lectin (as in next section) before cutting in order to evaluate choanocyte migration during the regenerative process.

### Cell dissociation-reaggregation experiments

Thousands of buds were collected in a Falcon tube (15mL) then slowly centrifuged, 10G 2min at RT. The supernatant of natural sea water (NSW) was discarded and the bud solution (1.5mL) was incubated in 20mL Calcium Magnesium Free Sea Water (CMSFW: 0.54 M NaCl, 10 mM KCl, 7 mM Na_2_SO_4_, 15 mM tris-HCl, 0.2mM NaHCO_3_, pH8) with 10mM EDTA pH8 and stirred for 1 h at RT. Cell suspension was filtered on 17 μm cell strainer. Concentration of cellular suspension was evaluated using C-Chip Neubauer. Before counting, the viability of cells, can be efficiently evaluated with fluorescein diacetate (FDA) and propidium iodide (PI) staining (Sigma) according to the protocol used by Sipkema et al. (2004).Then, 2mL of cell suspension (between 0.4.10^6^ and 0.56.10^6^ Cells/mL) were dispatched in 24 wells cell culture plate and CaCl_2_ solution was added to each well (final concentration 20mM). Culture plate was placed at 17°C and stirred very slowly on a 3D platform shaker. After reaggregation, cell aggregates were thus carefully transferred in 2mL Artificial Sea Water (ASW: 10mM CaCl_2_, 50mM MgCl2, 6H2O, 0.54M NaCl, 10 mM KCl, 7 mM Na_2_SO_4_, 15 mM tris-HCl, 0.2mM NaHCO_3_, pH 8) in 6 wells cell culture plates. Culture plates are placed at 17°C and stirred very slowly on a 3D platform shaker. The reaggregation process was monitored during 7 days. This manipulation has been repeated 3 times.

### Cell staining

Stock solution (1 mM) of the lipidic marker Cell Tracker™ CM-DiI Dye Red (553/570nm) (Molecular Probes) was prepared in dimethylsulfoxyde solution (DMSO). After short centrifugation, 2 μL were introduced in 1 mL NSW (1:500) and transferred in 24 wells cell plate with buds at 17°C ON.

All fluorescent (rhodamine or fluorescein labeled) lectins and agglutinins provided in vector laboratories kits (kit I (RLK-2200), kit II (FLK-3200) and kit III (FLK-4100)) were tested at different concentrations (1:100, 1:500, 1:1000, 1:2000 in NSW) for 15 min, 30min, 1 h at 17°C. After each treatment, buds were rinsed three times in NSW and immediately observed with an epifluorescence microscope DM2500 Leica and a confocal zeiss lsm 510 (for CM-Dil Dye treatment).

In a second step, the dilution and incubation times were optimized, for the 3 lectins that showed a more specific pattern: *Griffonia simplex* lectin 1 (Gsl1), *Phaseolus vulgaris Erythroagglutinin* (PhaE) and *Wheat germ agglutinin* (WGA).

For more precise observations samples were fixed with 3% PFA/PBS at 4°C ON, counterstaining with DAPI (1:500) (Thermo Fisher Scientific) and 1:1000 (iFluor 488 or Rhodamine or Alexa-647 (or 568) coupled-) phalloidin (Abcam, Thermo Fisher Scientific) and mounted in Prolong Diamond antifading mounting medium (Thermo Fisher Scientific); and then observed with confocal microscope Zeiss AxioImager Z2 780 or 880.

### Cell transfections

To develop a cell transfection protocol, we first used control nucleic acids molecules (BLOCKiT™ Fluorescent oligo control (Invitrogen™), pmaxGFP™ Vector (expressed under a CMV-promoter; LONZA) and a morpholino standard control oligo (Gene tools) with 3’ carboxyfluorescein green, in order to test transfection efficiency.

For soaking method, BLOCK-iT™ fluorescent oligo (20μM stock) were used at different final concentrations (2nM, 4nM, 8nM, 100nM) and pmaxGFP™ Vector (0.5μg/μL stock) at 5μg and 30μg in 5 mL natural sea water (NSW). Then, 3-5 buds were directly put in contact with these solutions placed of a 6 well cell culture plate and maintained at 17°C during 5H, 16H, 24H, 48H,72H, 5 days or 7 days without changing the sea water. The buds were rinsed 3 times with NSW and then observed with an epifluorescence microscope Leica DM2500 and a confocal microscope Zeiss after fixation and mounting.

For standard control morpholino (1mM stock) we tested different concentrations (10μM, 1μM, 0.9μM, 0,8μM, 0,6μM, 0,5μM and 0,1μm) in 500μL (filtered or not) NSW. The buds were placed in a 24 well cell plate (4 buds per well) and incubated at 17°C with the morpholino solutions during 24H, 48H, 72H, 96H and 120H. The buds were rinsed 1 time with NSW and fixed in PFA 3% O/N at 4°C. Alive buds were observed with an epifluorescence microscope Leica DM2500, and after counterstaining and mounting observed the samples with a confocal microscope Zeiss.

For testing the efficiency of lipofection for pmaxGFP™ Vector and BLOCK-iT™ fluorescent oligo, two lipofectamine transfection reagent (Invitrogen™), Lipofectamine^®^ 3000 and Lipofectamine^®^ LTX DNA (less cytotoxic) were tested in this study according the “24-well” protocol of the manufacturer’s instructions using 1μg DNA, but replacing the Opti-MEM™ medium, shown to be toxic for the buds by preliminary experiments, by PBS or filtered NSW. We used the indicated volume of DNA and reagent with each of the volume lipofectamine™ 3000 or LTX as preconized when performing optimization. Buds (N=24 for Lipofectamine^®^ 3000; N=60 for Lipofectamine^®^ LTX DNA for each of DNA constructs) were incubated at 17°C during 1 to 2 days and then rinsed with NSW before observation with an epifluorescence microscope DM2500 Leica. For morpholino standard control oligo, the different concentrations, as described above, were tested with aquaeous (mannitol-based) Endo Porter solution at 1μM, 2μM, 3μM and 6μM in 500μL NSW. The buds are treated in the same conditions as without EndoPorter (see above).

We also tested electroporation method for BLOCK-iT™ fluorescent oligo and pmaxGFP™ Vector introduction in *O. lobularis* cells using the Neon™ Transfection System and Neon™ Tip 100μL (Invitrogen). To avoid electric arcs, we replaced NSW by a Mannitol solution (0,77M) (Zeller et al., 2006). We used 6μg to 15μg of pmaxGFP™ Vector (0.5μg/μL stock) and 0.1 to 0.2μM of BLOCK-iT™ (20μM stock) for each electroporation. 14 buds were added to the Mannitol solution (120μL) with or without acids nucleic molecules and incubated 1 min. Then the mixture was transferred into tips and electroporated at different conditions (0 Volt-1mS-1 pulse / 500 volts-1mS-1 pulse /500volts-5mS-1pulse /500 volts-40mS-1pulse / 500 volts-1mS-5 pulses / 500V-10mS-2 pulses / 500 volts-5mS-1pulse / optimization of Neon system number 5 / 7 / 10 / 13/18/ 24). After electroporation, buds were placed in 6 mL NSW at 17°C (plate-6 wells) during 4, 24, 96, 120 or 168H before epifluorescence microscope observations.

## Acknowledgements

All the authors thank the imaging facilities of the France Bioimaging infrastructure and in particular Brice Detailleur, Laetitia Hudecek and Elsa Auroux-castellani for their help and advices to develop imaging in *O. lobularis* buds. We also acknowledge the diving facilities of the Institute OSU Pytheas (Frédéric Zuberer, Laurent Vanbostal and Dorian Guillemain) and divers from the IMBE lab (Anne Haguenauer, Pascal Mirleau) for *in situ* monitoring and collecting *O. lobularis*. We thank the molecular biology and morphology support services of IMBE for providing facilities needed to develop techniques. We also thank Elsa Bazellières for her help in launching bead monitoring and Jean Vacelet for helpful discussions.

We are grateful to Matthieu De La Rosa, Kassandra de Pao Mendonca, Damien Vary and Maxime Angely who participated in preliminary experimental assays during their internships.

